# Genome-wide screen for haploinsufficient cell size genes in the opportunistic yeast *Candida albicans*

**DOI:** 10.1101/084244

**Authors:** Julien Chaillot, Michael A Cook, Jacques Corbeil, Adnane Sellam

**Affiliations:** Infectious Diseases Research Centre (CRI), CHU de Québec Research Center (CHUQ), Université Laval, Quebec City, QC, Canada; Centre for Systems Biology, Samuel Lunenfeld Research Institute, Mount Sinai Hospital, Toronto, Canada M5G 1X5, Canada.; Department of Molecular Medicine, Université Laval, Québec, Canada; Department of Microbiology, Infectious Disease and Immunology, Faculty of Medicine, Université Laval, Quebec City, QC, Canada

**Keywords:** *Candida albicans*, cell size, haploinsufficiency, Start control

## Abstract

One of the most critical but still poorly understood aspects of eukaryotic cell proliferation is the basis for commitment to cell division in late G1 phase called Start in yeast and the Restriction Point in metazoans. In all species, a critical cell size threshold coordinates cell growth with cell division and thereby establishes a homeostatic cell size. While a comprehensive survey of cell size genetic determinism has been performed in the saprophytic yeasts *Saccharomyces cerevisiae* and *Schizosaccharomyces pombe*, very little is known in pathogenic fungi. As a number of critical Start regulators are haploinsufficient for cell size, we applied a quantitative analysis of the size phenome, using elutriation-barcode sequencing methodology, to 5,639 barcoded heterozygous deletion strains in the opportunistic yeast *Candida albicans*. Our screen identified conserved known regulators and biological processes required to maintain size homeostasis in addition to novel *C. albicans* specific size genes, and provided a conceptual framework for future mechanistic studies. Interestingly, some of the size genes identified were required for fungal pathogenicity suggesting that cell size homeostasis may be elemental to *C. albicans* fitness or virulence inside the host.

## INTRODUCTION

In eukaryotic species, growth and division are coupled at Start (Restriction point in metazoan), the point in late G1 at which the cell commits to the next round of division (Jorgensen and Tyers 2004). Cells must grow to reach a critical size threshold at Start and thereby establishes a homeostatic cell size. Pioneering studies in the eukaryotic model *Saccharomyces cerevisiae* revealed that a large proportion of the genome (>10%) and of cellular functions impact on the size of cells, in processes ranging from ribosome biogenesis (*Ribi*), mitochondrial function, to signal transduction and cell cycle control (Jorgensen et al. 2002; Zhang et al. 2002; Soifer and Barkai 2014). While follow-on studies revealed many crucial players in size regulation such as the G1 repressor Whi5 and the *Ribi* master regulators Sch9 and Sfp1, both the central mechanism by which cells sense their size and the means by which they alter their size set-point to meet environmental demands remain elusive (Turner et al. 2012).

*Candida albicans* is a diploid ascomycete yeast that is an important commensal and opportunistic pathogen in humans colonizing primarily mucosal surfaces, gastrointestinal and genitourinary tracts, and skin (Berman and Sudbery 2002). Interest in *C. albicans* is not limited to understanding its function as a pathogenic organism, as it has an ecological niche that is obviously distinct from the classic model ascomycete *S. cerevisiae. C. albicans* has served as an important evolutionary milepost with which to assess conservation of biological mechanisms. Recent investigations uncovered an extensive degree of rewiring of fundamental signalling and transcriptional regulatory networks as compared to *S. cerevisiae* and other fungi (Lavoie et al. 2009; Sellam et al. 2009; Blankenship et al. 2010; Homann et al. 2009; Sandai et al. 2012).

Haploinsufficiency is a phenotypic feature wherein a deletion of one allele in a diploid genome leads to a discernable phenotype. In eukaryotes, a number of critical size regulators such as the G1 cyclin Cln3 and the AGC kinase Sch9 in *S. cerevisiae*, and the Myc oncogene in *Drosphila melanogaster* are haploinsufficient (Sudbery et al. 1980) (Jorgensen et al. 2002; Barna et al. 2008). Here we exploited gene haploinsufficiency to identify genes and biological process that influence size control in *C. albicans*. Given the importance of *C. albicans* as an emerging eukaryotic model, very little is known regarding the genetic networks that control size homeostasis in this opportunistic yeast. A systematic screen using elutriation-based size fractioning (Cook et al. 2008) coupled to barcode-sequencing (Bar-seq) identified 685 genes (10 % of the genome) that influenced size control under optimal growth conditions. While *C. albicans* and *S. cerevisiae* share the morphological trait of budding, and core cell cycle and growth regulatory mechanisms (Berman 2006; Cote et al. 2009), a limited overlap was obtained when comparing the size phenome of both yeasts. This genome-wide survey will serve as a primary entry point into the global cellular network that couples cell growth and division in *C. albicans*.

## MATERIALS AND METHODS

### Strains and growth conditions

*C. albicans* SC5314 and CAI4 (*ura3*::imm434/*ura3*::imm434 *iro1/iro1*::imm434) (Fonzi and Irwin 1993) wild-type (WT) strains, and mutants of the Merck DBC (Double BarCoded) heterozygous diploid collection (Xu et al. 2007) were routinely maintained at 30°C on YPD (1% yeast extract, 2% peptone, 2% dextrose, with 50 mg/ml uridine) or synthetic complete (SC; 0.67% yeast nitrogen base with ammonium sulfate, 2.0% glucose, and 0.079% complete supplement mixture) media. The Merck DBC collection is available for public distribution through the NRC’s Royalmount Avenue research facility (Montreal, Canada).

### Combination of *C. albicans* mutants into a single pool

A sterilized 384-well pin tool was used to transfer DBC mutant cells into Nunc Omni Trays containing YPD-agar and colonies were grown for 48 hours at 30°C. Missing or slow growing colonies were grown separately by repinning 3.5 µl from the initial liquid cultures. Each plate was overlaid with 5 ml of YPD and cells were resuspended using Lazy-L spreader and harvested by centrifugation for 5 min at 1800 g. The obtained cell pellet was resuspended in 20 ml fresh YPD and DMSO was added to 7% (vol/vol). Mutant pools were aliquoted and stored at −80°C.

### Cell size selection by centrifugal elutriation

The mutant pool was size-fractioned using centrifugal elutriation with the Beckman JE-5.0 elutriation system. This technique separates cells on the basis of size. A tube of pooled mutant population was thawed on ice and used to inoculate 2 L of YPD at an OD_595_ of 0.05. Mutant cells were grown for four generations at 30°C under agitation to reach approximatively 5 x10^10^ cells.

Cells were then pelleted by centrifugation and resuspended in 50 ml fresh YPD. To disrupt potential cell clumps and separate weakly attached mother and daughter cells, the 50 ml pooled cells were gently sonicated twice for 30s. The resuspended cells were directly loaded into the elutriator chamber of the Beckman JE-5.0 elutriation rotor. A 1 ml sample of cells was retained separately as a pre-elutriated cell fraction. The flow rate of the pump was set to 8 mL/min to ensure the loading of cells.

To elute small cell size mutant fractions, the pump flow rate was increased in a step-wise fashion (in 2-4 mL/min increments). For each flow rate, a volume of 250 ml was collected from the output line of the rotor.

### Barcode sequencing

Barcode sequencing (Bar-seq) was performed using Illumina HiSeq2500 platform. Genomic DNA was extracted from each cell fraction using YeaStar kit (Zymo Research). The 20-bp UpTag barcode of each strain were amplified by PCR (Xu et al. 2007). Primers used for PCR recognize the common region of each barcode, contain the multiplexing tag and sequences required for hybridization to the Illumina flowcell. PCR products were purified from an agarose gel using the QIAquick Gel Extraction kit (Qiagen) and quantified by QuantiFluor^®^ dsDNA System (Promega). Bar-seq data was processed as following: after filtering out low frequency barcode counts, the complete set of replicate barcode reads were normalized using a cyclic loess algorithm (R package “limma”). Reads from individual elutriation fractions, relative to the pre-elutriation population, were further M-A loess normalized and converted to Z scores.

### Confirmation of cell size phenotypes

Cell size determination was performed using a Z2-Coulter Counter channelizer (Beckman). *C. albicans* cells were grown overnight in YPD at 30°C, diluted 1000-fold into fresh YPD and grown for 5h at 30°C to reach a final density of 5. 10^6^-10^7^ cells/ml, a range in which size distributions of the different WT strain used in this study do not change. A total of 100 µl of exponentially growing cells was diluted in 10 ml of Isoton II electrolyte solution, sonicated three times for 10s and used for size quantification in the Z2-Coulter. Size distribution data were normalized to cell counts in each of 256 size bins. Data analysis and size distribution clustering were performed using custom R scripts.

### Determination of critical cell size

Critical sizes of *cln3/CLN3, cdc28/CDC28* and *sch9/SCH9* mutants were determined using budding index as a function of size. G1 daughter cells were obtained using the JE-5.0 centrifugal elutriation system (Beckman) as described previously (Tyers et al. 1993). *C. albicans* G1-cells were released in fresh YPD medium and fractions were harvested at an interval of 10 min to monitor bud index. Additional fractions were collected to assess transcript levels of the *RNR1* and *ACT1* as cells progressed along G1 phase.

### Real-time quantitative PCR

A total of 10^8^ G1 phase cells were harvested, released into fresh YPD medium and grown for 10 min prior to harvesting by centrifugation and stored at −80°C. Total RNA was extracted using the RNAeasy purification kit (Qiagen) and glass bead lysis in a Biospec Mini 24 bead-beater as previously described (Sellam et al. 2009). cDNA was synthesized from 2 µg of total RNA using the SuperScript^®^ III Reverse Transcription system (50 mm Tris-HCl, 75 mm KCl, 10 mm dithiothreitol (DTT), 3 mm MgCl2, 400 nm oligo(dT)15, 1 m random octamers, 0.5 mm dNTPs, and 200 U Superscript III reverse transcriptase). The total volume was adjusted to 20 µl, and the mixture was then incubated for 60 min at 42°C. Aliquots of the resulting first-strand cDNA were used for real-time qPCR amplification experiments. qPCR was performed using the iQ^™^ 96- well PCR system for 40 amplification cycles and QuantiTect SYBR Green PCR master mix (Qiagen). Transcript levels of *RNR1* were estimated using the comparative Ct method as described by Guillemette *et al.* (Guillemette et al. 2004) and the *C. albicans ACT1* ORF as a reference. The primer sequences were: RNR1-forward: 5’- GACTATCTACCATGCTGCTGTTG-3’; RNR1-reverse: 5’- GGTGCAACCAACAAGGAGTT-3’; ACT1-forward 5’-GAAGCCCAATCC AAAAGA-3’ and ACT1-reverse 5’-CTTCTGGAGCAACTCTCAATTC-3’.

### Gene ontology analysis

Gene ontology (GO) term enrichment of size mutants was determined using the Generic GO Term Finder tool (http://go.princeton.edu/cgi-bin/GOTermFinder) with multiple hypothesis correction (Boyle et al. 2004). Descriptions related to gene function in **Table S1** were extracted from CGD (Candida Genome Database) database (Inglis et al. 2012). Information related to gene essentiality/dispensability was taken from O’Meara *et al.* (O’Meara et al. 2015) and CGD database.

## RESULTS AND DISCUSSION 3

### The Cln3-Cdc28 kinase complex and Sch9 are haploinsufficient for cell size

In *S. cerevisiae*, a number of critical Start regulators are haploinsufficient for cell size, including the rate-limiting G1 cyclin Cln3 and a number of essential ribosome biogenesis factors such as the the AGC kinase Sch9 (Sudbery et al. 1980; Jorgensen et al. 2002). To test whether size haploinsufficiency exists in *C. albicans* homologs, size distributions of the AGC kinase Sch9, the cyclin Cln3 G1 and its associated cyclin-dependant kinase Cdc28 heterozygous mutants were examined. Both *cln3/CLN3* and *cdc28/CDC28* showed an increase of size as compared to their congenic parental strain with a median sizes 13 % (59 fL) and 19 % (62 fL) larger than the WT strain (52 fL), respectively (Figure 1A). As in *S. cerevisiae, sch9/SCH9* exhibited a reduced size of about 23 % (40 fL) as compared to WT.

**Figure 1.**
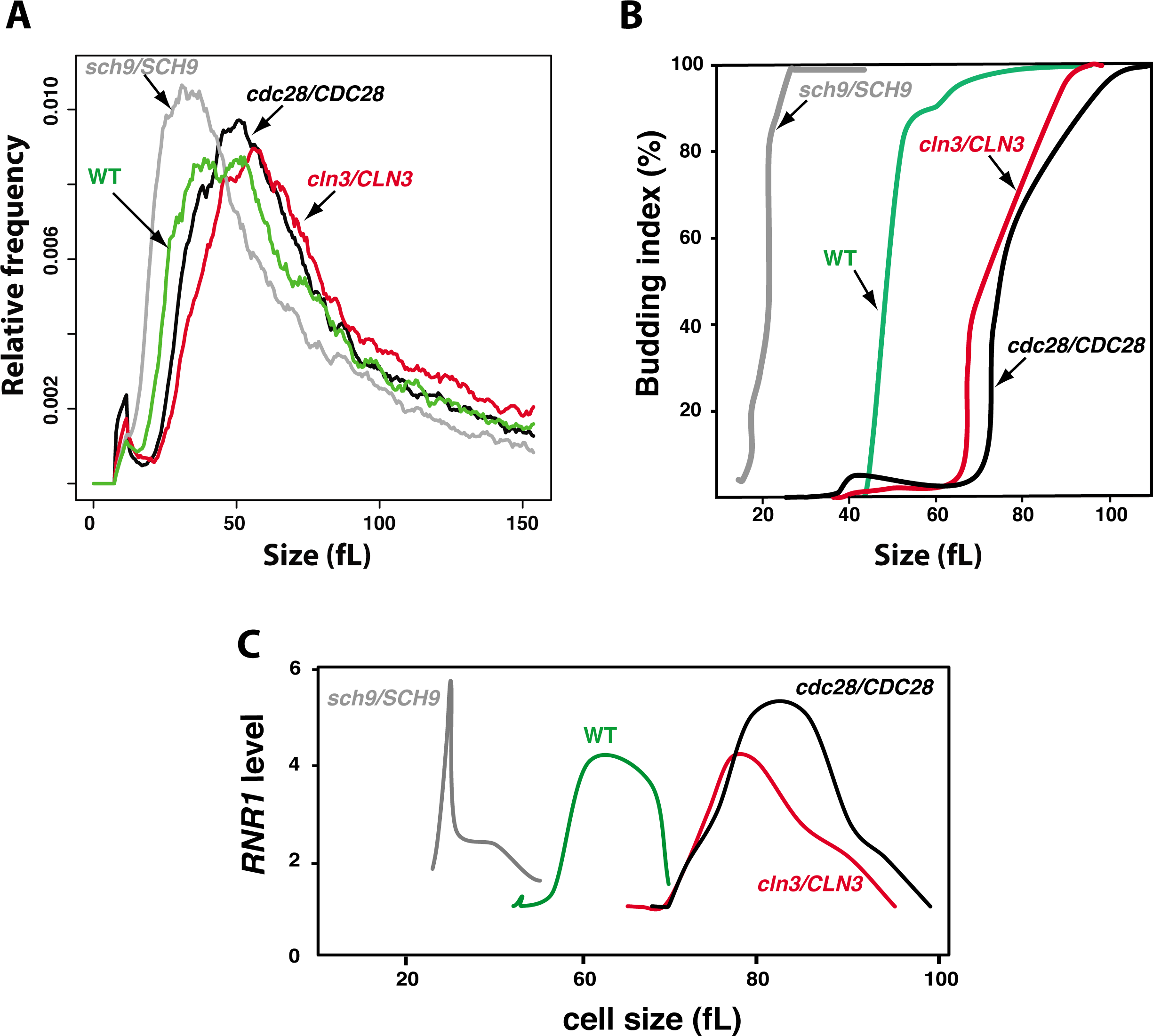
The Cln3-Cdc28 kinase complex and the AGC kinase Sch9 control Start in *C. albicans.* **(A)** Size distributions of the WT strain (CAI4) as compared to *lge* mutants *cln3/CLN3* and *cdc28/CDC28*, as well as the *whi* mutant *sch9/SCH9*. (**B-C**) Start is delayed in *cln3/CLN3* and *cdc28/CDC28* and accelerated in *sch9/SCH9.* (**B**) Elutriated G1 phase daughter cells were released into fresh media and monitored for bud emergence as a function of size. (**C**) G1/S transcription. *RNR1* transcript level was assessed by quantitative real-time PCR and normalized to *ACT1* levels.

Two hallmarks of Start, namely SBF-dependent transcription and bud emergence, were delayed in both *cln3/CLN3* and *cdc28/CDC28* and accelerated in *sch9/SCH9* demonstrating that the Cln3-Cdc28 complex and Sch9 regulate the cell size threshold at Start. The *cln3/CLN3* mutant passed Start after growing to 92 fL, 24 % higher than the parental WT cells, which budded at 74 fL (Figure 1B). Similarly, *cdc28/CDC28* reached Start at 105 fL which is 41 % higher than WT. The onset of G1/S transcription was delayed in both mutants as judged by the expression peak of the G1-transcript *RNR1* (Figure 1C). The small mutant *sch9/SCH9* passed start at 30 fL, a size 60 % less than the WT and displayed accelerated G1/S transcription (Figure 1B-C). These data demonstrate that, as in *S. cerevisiae*, size haploinsufficiency in *C. albicans* can be used to screen for dosage dependent regulators of growth and division at Start.

### A high-throughput screen for cell size haploinsufficiency

To identify all dosage-sensitive regulators of size in *C. albicans*, a genome-wide screen was performed where pooled mutants were separated based on their size by centrifugal elutriation and their abundance determined by Bar-seq. This method has been previously validated in *S. cerevisiae* (Cook et al. 2008), yielding a high degree of overlap when compared to a strain-by-strain analyses (Jorgensen et al. 2002) (Figure 2A). In the current study, we screened a comprehensive set of 5470 heterozygous deletion diploid strains from the Merck DBC collection (Xu et al. 2007) for cell size defects. This collection covers 90 % of the 6046 protein coding open reading frames based on the current CGD annotation (Binkley et al. 2014). Two small cell size fractions were obtained by centrifugal elutriation and were used for these experiments (Figure 2B). Small cells and corresponding small deletion mutants are enriched in these fractions, while large cells strains are depleted. To determine mutant abundance in each fraction, genomic DNA of each pool was extracted and barcodes were PCR-amplified and sequenced. Abundance of each mutant in each fraction was appreciated by calculating the ratio of elutriated cells counts over counts of pre-elutriated cells.

**Figure 2.**
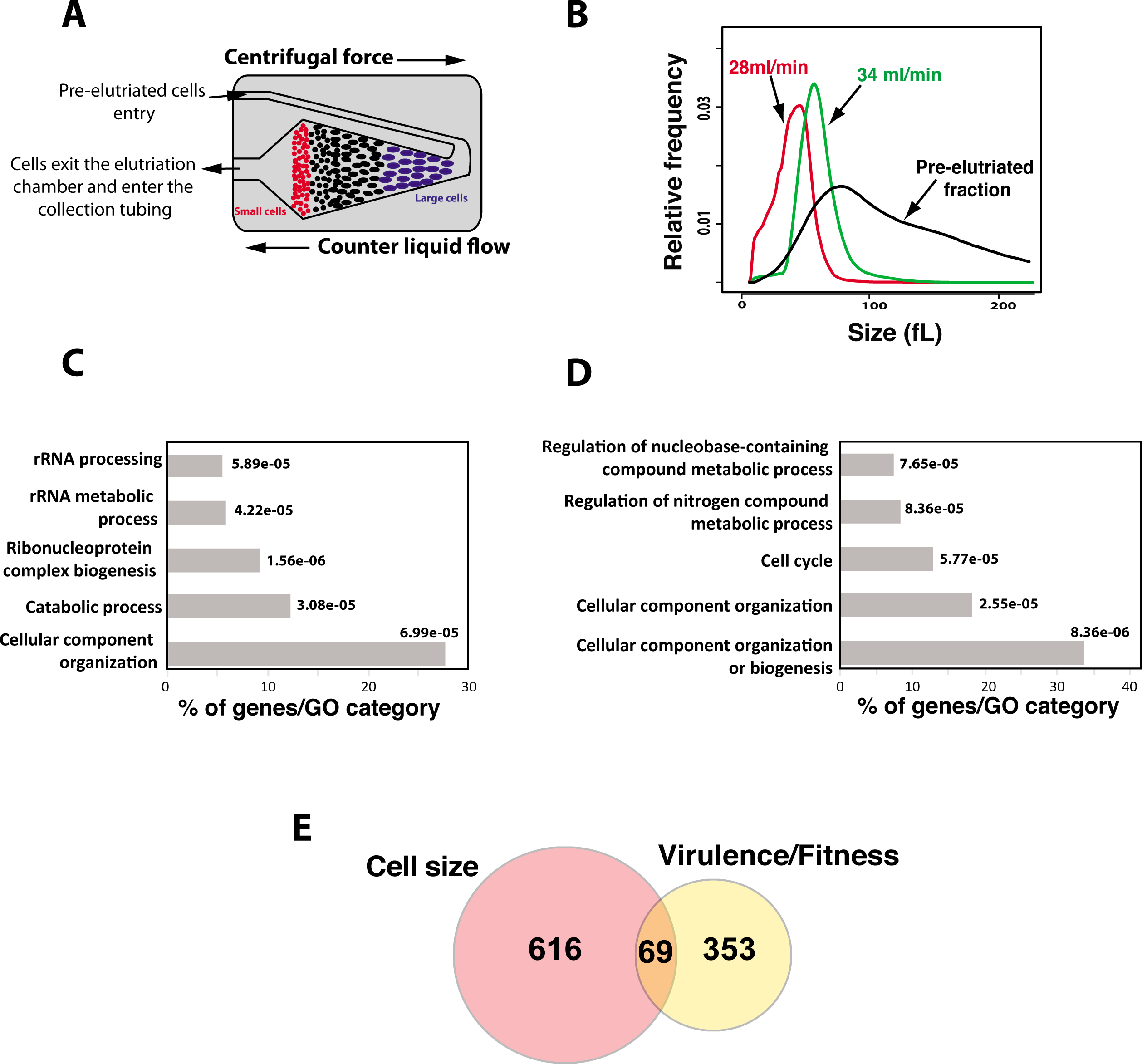
Systematic cell size screen using molecular barcode-elutriation and Bar-seq. **(A)** Centrifugal elutriation separates cells on the basis of size. Progressive increase of the rate of flow of liquid medium counter to the direction of centrifugal force elutes yeast cells of increasingly larger size from the chamber. (**B**) Size distributions of pooled DBC mutants before (pre-elutriated) and after elutriation of two small cell fractions (28 and 34 ml/min). (**C-D**) GO terms enrichment of *whi* (**C**) and *lge* (**D**) mutants (p > 1e-05). GO analysis was performed using GOTermFinder (http://go.princeton.edu/cgi-bin/GOTermFinder). (**E**) Overlap between *C. albicans* genes haploinsufficient for cell size and those affecting virulence phenotypes. Avirulent mutant phenotypes were obtained from CGD based on decreased competitive fitness in mice and/or reduced invasion and damage to host cells.

To identify mutants with size defects, a two-step filter was applied. First, a size cut-off value was determined based on a benchmark set of conserved small (*sch9/SCH9*) and large (*cln3/CLN3 and cdc28/CDC28*) sized mutants for which size was reduced or increased at least 12 % as compared to the parental WT strain. Second, a normalized z-score of 1.5 and −1.5 was used to identify both small (*whi*) and large (*lge*) size mutants, respectively. A total of 12 size mutants were excluded from our analysis since they were found in both *whi* and *lge* datasets. Microscopic examination revealed that these mutants grew predominantly as pseudohyphae. Based on these criteria, we identified 685 mutants that exhibited a size defect in both elutriated fractions. This includes a 382 *whi* and 303 *lge* mutants (**Table S1**). As expected, *cln3/CLN3* and *cdc28/CDC28* mutants were identified as *lge* mutants while *sch9/SCH9* was found among the smallest mutant in the elutriated pools. A total of 15 *whi* and 15 *lge* mutants were randomly selected and their size was measured by electrolyte displacement on a Coulter Z2 channelizer. The obtained data confirmed size defect in all 40 mutants examined (**Table S2**).

### Synthesis of ribosome and cell cycle are required for cell size homeostasis

Gene Ontology (GO) enrichment analysis revealed that mutation in genes related to rRNA processing and ribosome biogenesis confer small cell size, while mutations of cell cycle genes resulted in *lge* phenotype (Figure 2C-D and Table S1). Heterozygous deletion of genes of different functional categories related to protein translation including rRNA processing (*CSL4, UTP7, DIS3, NOP53, FCF2, UTP23, DIP2, UTP15, SAS10*), ribosome exports (*RIX7, RRS1, NUP84, NUP42, RPS5, NOG1*), translation elongation (*RIA1, EFT2, CEF3*) and transcription of RNA Pol I and III promoters (*CDC73, RPB8, RPA49, RPB10, SPT5, RPA12, RPC25*) exhibited a *whi* phenotype. Heterozygous deletion mutation of structural components of both cytoplasmic (*RPL18, RPL20B, RPL21A, RPS5, UBI3*) and mitochondrial (*RSM24, RSM26, NAM9, MRPL20*) ribosomes decreased cell size. As in *S. cerevisiae*, mutants of the ribosome biogenesis regulator Sch9 and the transcription factor Sfp1 had a small size. Overall, as shown in other eukaryotic organisms, these data lead to the hypothesis that the rate of ribosome biogenesis or translation is a critical element underlying cell size control (Jorgensen et al. 2004). Haploinsufficient *whi* mutants corresponded also to catabolic processes associated mainly with ubiquitin-dependent proteolysis (*DOA1, GRR1, UBP1, UBP2, UFD2, SSH4, UBX7, RPN2, TIP120, TUL1, RPT2, PRE1, GID7*).

*Lge* mutants were predominantly defective in functions related to the mitotic cell cycle (Figure 2D and Table S1). These mutants include genes required for G1/S transition (G1 cyclin Cln3 and Ccn1, Cdc28 and Met30) suggesting that delay in G1 phase is the primary cause of their increased size. We also found that mutations in processes related to DNA replication (*ORC3, ORC4, MCM3, CDC54, RFC3, PIF1, SMC4, ELG1*), G2/M transitions (Hsl1, Cdc34) and cytoskeleton-dependant cytokinesis (*MYO5, INN1, SEC15, CDC5, CHS1*) conferred an increase of cell size. These data suggest that cell size homeostasis could be controlled in all cell cycle phases. A similar observation was reported in *S. cerevisiae* where a genome-wide microscopic quantitative size survey uncovered that mutants of the G2/M transition and mitotic exit failed to properly control their size (Soifer and Barkai 2014). Other investigations support this hypothesis and propose a model where, in addition to the G1-phase, size is sensed and controlled at G2/M checkpoint (Anastasia et al. 2012; King et al. 2013; Harvey and Kellogg 2003).

While a large proportion of *whi* mutants in *C. albicans* were related to ribosome biogenesis, inactivation of genes controlling translation initiation (*ASC1, SCD6, PAB1, GCD6, GCD2, SUI1, EIF4E, GCD11*) resulted in *lge* phenotype. A similar finding was reported in different genome-scale surveys of size phenome in *S. cerevisiae* (Jorgensen et al. 2002; Soifer and Barkai 2014). This large size phenotype in these mutants could be explained by the fact that regulators of Start onset, such as G1-cyclins Cln3 (Barbet et al. 1996; Polymenis and Schmidt 1997), are sensitive to the rate of translation initiation.

### Plasticity of size phenome and *C. albicans* fitness

Recent evidence has uncovered an extensive degree of rewiring of both cis-transcriptional regulatory circuits and signalling pathways across many cellular and metabolic processes between the two budding yeasts, *C. albicans* and *S. cerevisiae* (Lavoie et al. 2010; Li and Johnson 2010; Blankenship et al. 2010; Lavoie et al. 2009). In *S. cerevisiae*, a similar size haploinsufficiency screen was performed in heterozygous diploid strains of essential genes (Jorgensen et al. 2002). To assess the extent of conservation and plasticity of the size phenome, genes that were haploinsufficient for cell size in *C. albicans* were compared to their corresponding orthologs in *S. cerevisiae*. This analysis revealed a limited overlap between the two species with five *whi* (*rpl18a, sch9, rlp24, nop2, nog1*) and two *lge* (*rpt4, cln3*) mutants in common. In fact, genes with reciprocal size phenotypes were similar in frequency (the *whi* mutants *rpt2/RPT2* and *pkc1*\*PKC1* in *C. albicans* had *lge* phenotype in *S. cerevisiae*).

Interestingly, the corresponding homozygous deletion mutants of many *C. albicans* haploinsufficient size genes were shown to be required for virulence. A total of 69 size genes (representing ~ 10%), including 47 small and 22 large size mutants, in our dataset were linked to *C. albicans* virulence or adaptation in the human host (Figure 2E). This suggests that cell size is an important virulence trait that can be targeted by antifungal therapy. Hypothetically, virulence defect in small size mutant could be linked to the reduced surface of the contact interface between *C. albicans*, with either host cells or medical devices in case of biofilm infections. Indeed, we have previously shown that the *whi* transcription factor mutant *ahr1* had attenuated virulence and exhibited a decreased attachment ability to abiotic surface such polystyrene, which consequently impaired biofilm formation (Askew et al. 2011). On the other hand, virulence defect in *lge* mutant could be associated with the fact that cells with large surfaces had a decreased lifespan which might impact their fitness and their viability inside the host (Yang et al. 2011; Mortimer and Johnston 1959).

While the link between *C. albicans* size and virulence remains uncharacterized, many investigations reported that many other fungal pathogens such as *Cryptococcus neoformans* and *Mucor circinelloides* adjust their cell size to access to specific niche in the host or to escape from immune cells (Wang and Lin 2012). In *C. albicans*, recent investigations have shown that large GUT (Gastrointestinallly Induced Transition) cells, as compared to the standard yeast form, define the commensal form of this fungus (Pande et al. 2013). Furthermore, Tao *et al.* (Tao et al. 2014) recently uncovered a novel intermediate phase between the white and the *C. albicans* mating competent opaque phenotypes called the Gray phenotype. The Gray cells are similar to opaque cells in general shape, however, they exhibit a small size and low mating efficiency. The Gray cell type has unique virulence characteristics, with a high ability to cause cutaneous infections and a reduced capacity in colonizing internal organs such as kidney, lung and brain. Taken together, these lines of evidence emphasize the possible link between cell size and *C. albicans* fitness.

In summary, we provided the first comprehensive genome-wide survey of haploinsufficient cell size in a eukaryotic organism. In contrast to the case of *S. cerevisiae*, where a similar screen was limited to essential genes (Jorgensen et al. 2002), our screen spanned the genome. A total of 300 (43.8 %) dispensable genes and only 87 (12.7 %) of essential genes were haploinsufficient for size. Overall, our screen identified known conserved regulators (Sch9, Sfp1, Cln3) and biological processes (ribosome biogenesis and cell cycle control) required to maintain size homeostasis in this opportunistic yeast. We also identified novel *C. albicans* size specific genes and provided a conceptual framework for future mechanistic studies. Interestingly, some of the size genes identified were required for fungal pathogenicity suggesting that cell size homeostasis may be elemental to *C. albicans* fitness or virulence inside the host.

## ACKNOWLEDGMENTS

Work in Adnane Sellam’s laboratory is supported by Fonds de Recherche du Québec-Santé (FRQS) (Établissement de jeunes chercheurs) and the Natural Sciences and Engineering Research Council of Canada (NSERC) Discovery Grant (06625). Adnane Sellam is a recipient of the Fonds de Recherche du Québec-Santé (FRQS) J1 salary award. We thank Lynda Robitaille for technical assistance and NRC Canada for providing the Merck DBC mutants used in this work.

## Supplemental Tables

**Table S1.** List of *whi* and *lge* haploinsufficient size mutants in *C. albicans* grouped according to GO biological process terms.

**Table S2.** Confirmation of cell size defect in 15 *whi* and 15 *lge* mutants using the Z2-Coulter counter channelizer.

## LITERATURE CITED

Anastasia, S.D., D.L. Nguyen, V. Thai, M. Meloy, T. MacDonough et al., 2012 A link between mitotic entry and membrane growth suggests a novel model for cell size control. J Cell Biol 197 (1):89–104.

Askew, C., A. Sellam, E. Epp, J. Mallick, H. Hogues et al., 2011 The zinc cluster transcription factor Ahr1p directs Mcm1p regulation of *Candida albicans* adhesion. Mol Microbiol 79:940–953.

Barbet, N.C.U., Schneider, S.B. Helliwell, I. Stansfield, M.F. Tuite et al., 1996 TOR controls translation initiation and early G1 progression in yeast. Mol Biol Cell 7 (1):25–42.

Barna, M., A. Pusic, O. Zollo, M. Costa, N. Kondrashov et al., 2008 Suppression of Myc oncogenic activity by ribosomal protein haploinsufficiency. Nature 456 (7224):971–975.

Berman, J., 2006 Morphogenesis and cell cycle progression in Candida albicans. Curr Opin Microbiol 9 (6):595–601.

Berman, J., and P.E. Sudbery, 2002 Candida Albicans: a molecular revolution built on lessons from budding yeast. Nat Rev Genet 3 (12):918–930.

Binkley, J., M.B. Arnaud, D.O. Inglis, M.S. Skrzypek, P. Shah et al., 2014 The Candida Genome Database: the new homology information page highlights protein similarity and phylogeny. Nucleic Acids Res 42 (Database issue):D711–716.

Blankenship, J.R., S., Fanning, J.J., Hamaker, and A.P. Mitchell, 2010 An extensive circuitry for cell wall regulation in Candida albicans. PLoS Pathog 6 (2):e1000752.

Boyle, E.I., S., Weng, J., Gollub, H., Jin, D. Botstein et al., 2004 GO::TermFinder–open source software for accessing Gene Ontology information and finding significantly enriched Gene Ontology terms associated with a list of genes. Bioinformatics 20 (18):3710–3715.

Cook, M.A., C.K., Chan, P. Jorgensen, T. Ketela, D. So et al., 2008 Systematic validation and atomic force microscopy of non-covalent short oligonucleotide barcode microarrays. PLoS One 3 (2):e1546.

Cote, P., H. Hogues, and M. Whiteway, 2009 Transcriptional analysis of the Candida albicans cell cycle. Mol Biol Cell 20 (14):3363–3373.

Fonzi, W.A., and M.Y. Irwin, 1993 Isogenic strain construction and gene mapping in Candida albicans. Genetics 134 (3):717–728.

Guillemette, T., A. Sellam, and P. Simoneau, 2004 Analysis of a nonribosomal peptide synthetase gene from Alternaria brassicae and flanking genomic sequences. Curr Genet 45 (4):214–224.

Harvey, S.L., and D.R. Kellogg, 2003 Conservation of mechanisms controlling entry into mitosis: budding yeast wee1 delays entry into mitosis and is required for cell size control. Curr Biol 13 (4):264–275.

Homann, O.R., J., Dea, S.M., Noble, and A.D. Johnson, 2009 A phenotypic profile of the Candida albicans regulatory network. PLoS Genet 5 (12):e1000783.

Inglis, D.O., M.B., Arnaud, J. Binkley, P. Shah, M.S. Skrzypek et al., 2012 The Candida genome database incorporates multiple Candida species: multispecies search and analysis tools with curated gene and protein information for Candida albicans and Candida glabrata. Nucleic Acids Res 40 (Database issue):D667–674.

Jorgensen, P., J.L. Nishikawa, B.J. Breitkreutz, and M. Tyers, 2002 Systematic identification of pathways that couple cell growth and division in yeast. Science 297 (5580):395–400.

Jorgensen, P., I. Rupes, J.R. Sharom, L., Schneper, J.R. Broach et al., 2004 A dynamic transcriptional network communicates growth potential to ribosome synthesis and critical cell size. Genes Dev 18 (20):2491–2505.

Jorgensen, P., and M., Tyers, 2004 How cells coordinate growth and division. Curr Biol 14 (23):R1014–1027.

King, K., H. Kang, M. Jin, and D.J. Lew, 2013 Feedback control of Swe1p degradation in the yeast morphogenesis checkpoint. Mol Biol Cell 24 (7):914–922.

Lavoie, H., H. Hogues, J. Mallick, A. Sellam, A. Nantel et al., 2010 Evolutionary tinkering with conserved components of a transcriptional regulatory network. PLoS Biol 8 (3):e1000329.

Lavoie, H., H. Hogues, and M. Whiteway, 2009 Rearrangements of the transcriptional regulatory networks of metabolic pathways in fungi. Curr Opin Microbiol 12 (6):655–663.

Li, H., and A.D. Johnson, 2010 Evolution of transcription networks–lessons from yeasts. Curr Biol 20 (17):R746–753.

Mortimer, R.K., and J.R. Johnston, 1959 Life span of individual yeast cells. Nature 183 (4677):1751–1752.

O’Meara, T.R., A.O. Veri, T. Ketela, B. Jiang, T. Roemer et al., 2015 Global analysis of fungal morphology exposes mechanisms of host cell escape. Nat Commun 6:6741.

Pande, K., C. Chen, and S.M. Noble, 2013 Passage through the mammalian gut triggers a phenotypic switch that promotes Candida albicans commensalism. Nat Genet 45 (9):1088–1091.

Polymenis, M., and E.V. Schmidt, 1997 Coupling of cell division to cell growth by translational control of the G1 cyclin CLN3 in yeast. Genes Dev 11 (19):2522–2531.

Sandai, D., Z. Yin, L. Selway, D. Stead, J. Walker et al., 2012 The evolutionary rewiring of ubiquitination targets has reprogrammed the regulation of carbon assimilation in the pathogenic yeast Candida albicans. MBio 3 (6).

Sellam, A., F. Tebbji, and A. Nantel, 2009 Role of Ndt80p in sterol metabolism regulation and azole resistance in Candida albicans. Eukaryot Cell 8 (8):1174–1183.

Soifer, I., and N., Barkai, 2014 Systematic identification of cell size regulators in budding yeast. Mol Syst Biol 10:761.

Sudbery, P.E., A.R., Goodey, and B.L. Carter, 1980 Genes which control cell proliferation in the yeast Saccharomyces cerevisiae. Nature 288 (5789):401–404.

Tao, L., H. Du, G. Guan, Y. Dai, C.J. Nobile et al., 2014 Discovery of a “white-gray-opaque” tristable phenotypic switching system in candida albicans: roles of non-genetic diversity in host adaptation. PLoS Biol 12 (4):e1001830.

Turner, J.J., J.C., Ewald, and J.M. Skotheim, 2012 Cell size control in yeast. Curr Biol 22 (9):R350–359.

Tyers, M., G. Tokiwa, and B. Futcher, 1993 Comparison of the Saccharomyces cerevisiae G1 cyclins: Cln3 may be an upstream activator of Cln1, Cln2 and other cyclins. EMBO J 12 (5):1955–1968.

Wang, L., and X., Lin, 2012 Morphogenesis in fungal pathogenicity: shape, size, and surface. PLoS Pathog 8 (12):e1003027.

Xu, D., B. Jiang, T. Ketela, S. Lemieux, K. Veillette et al., 2007 Genome-wide fitness test and mechanism-of-action studies of inhibitory compounds in Candida albicans. PLoS Pathog 3 (6):e92.

Yang, J., H. Dungrawala, H. Hua, A. Manukyan, L. Abraham et al., 2011 Cell size and growth rate are major determinants of replicative lifespan. Cell Cycle 10 (1):144–155.

Zhang, J., C. Schneider, L. Ottmers, R. Rodriguez, A. Day et al., 2002 Genomic scale mutant hunt identifies cell size homeostasis genes in S. cerevisiae. Curr Biol 12 (23):1992–2001.

